# pHAPE: a plasmid for production of DNA size marker ladders for gel electrophoresis

**DOI:** 10.1101/2022.11.04.515137

**Authors:** Angel G Allen, Karissa Barthelson, Michael Lardelli

## Abstract

DNA size markers (also known as “molecular weight markers” or “DNA ladders”) are an essential tool when using gel electrophoresis to identify and purify nucleic acids. However, the cost of these DNA ladders is not insignificant and, over time, impinges on the funds available for research and training in molecular biology. Here, we describe a method for the generation of “pHAPE”, a plasmid from which a variety of DNA ladders can be generated via simple restriction enzyme digestions. The pHAPE plasmid can be generated by mutagenesis of the commonly used pBluescript II SK+ phagemid followed by insertion of a custom 7,141 bp sequence (made up of three smaller fragments). Our use of pHAPE allows us some small relief from the ever-rising costs of performing molecular biology experiments (“Don’t worry, pHAPE”).

## 1 Introduction

Gel electrophoresis for the identification and purification of nucleic acids is a fundamental technique in molecular biology. Double-stranded DNA moves through electrophoresis gels at rates inversely related to fragment length/size (Helling et al., 1974), allowing the lengths of DNA fragments to be estimated by comparison to markers of known length (collections of which are commonly referred to as “size markers”, “molecular weight markers” or “DNA ladders”).

The first source of molecular weight markers that found widespread use was DNA from the lambda bacteriophage digested with restriction enzymes such as *Eco*RI or *Hind*III (Maniatis, 1982). This produces a versatile but irregular range of DNA fragment lengths. Inconveniently, the use of the bacteriophage lambda fragments to estimate the length of other DNA fragments of interest often required the measurement of electrophoresis migration distances and comparison of these using log-linear plots (Helling et al., 1974). Consequently, the commercial provision of DNA ladders with convenient fragment length intervals and different scales (e.g. 1 kb increments up to ~10 kb, 100 bp increments up to 1 kb) proved very popular and the regular purchase of such ladders is now common practice. However, the cost of these DNA ladders is not insignificant and, over time, impinges on the funds available for research and training in molecular biology.

The drive to minimise costs in molecular biology has encouraged laboratories to find methods to generate their own size markers, although all have their caveats. The cost of lambda phage DNA has been reported to be increasing in addition to it being harder to source (Henrici et al., 2017). A number of PCR-based techniques for ladder production have been described, although these are generally more suited to specific ladder designs (with fewer bands) and the materials required can be costly (Abbasian et al., 2015; Gopalakrishnan et al., 2010; Mostaan et al., 2015; Zhang et al., 2012). Progress in DNA synthesis technology has allowed the design and construction of plasmids specifically for the generation of DNA ladders after cleavage with restriction enzymes, for example, the pPSU plasmids produced by Henrici et al. (2017). These are attractive sources of DNA size markers as plasmids are easy to propagate at low cost, and molecular biology laboratories hold restriction enzymes as standard tools. Unfortunately, intellectual property (IP) constraints can hinder the distribution and use of these plasmids.

To bypass IP issues we embarked on a student project to design, assemble, and test a size marker plasmid. The plasmid is based on the pBluescript II SK+ phagemid that is widely available and in common use (Short et al., 1988). The plasmid can easily be recreated by synthesis of the described insert sequences and their ligation into a modified form of pBluescript II SK+. We named the resulting plasmid pHAPE (after the initials of the restriction enzymes required to produce its largest ladder output).

Multiple size markers can be produced from restriction enzyme digestion of pHAPE including a ladder featuring 1 kb increments (and with fragments ranging from 100 bp to 10 kb) and a ladder with 50 bp and 100 bp increments (spanning 50 bp to 1.2 kb). Our use of pHAPE allows us to further reduce molecular biology consumables costs (“Don’t worry, pHAPE”).

## 2 Results and Discussion

### 2.1 Design considerations

The pHAPE plasmid is based on the pBluescript II SK+ phagemid, a high copy number vector conferring ampicillin resistance (Short et al., 1988). To give the plasmid a total size of exactly 10,000 bp, a 7,141 bp insert based on a random sequence generated by the FaBox online toolbox (Villesen, 2007) constrained to contain equal quantities of A, C, G & T nucleotide residues is cloned into the *Kpn*I and *Sac*I sites of a mutated form of this vector. We designed the insert sequence with the intention that it should possess no evolved or designed function in cells. The random sequence generated initially is then modified to possess the desired restriction enzyme recognition sequences only at appropriate positions. The final sequence that is inserted into a modified form of pBluescript II SK+ to construct pHAPE can be found in the Supplementary Data File.

We desired 2 ladders encompassing different fragment length ranges: 1) A versatile “HAPE ladder”, with fragments ranging from 10,000 bp down to 100 bp and including 100 bp increments between 1 kb and 100 bp. 2) A ladder including 50-100 bp increments, specifically for use on high percentage agarose gels for estimating the lengths of low molecular weight DNA fragments of less than 1 kb (now referred to as the “*Bam*HI ladder” Table 1). An additional design consideration is that the visibility of stained DNA “bands” in electrophoresis gels varies with the total DNA mass in the band. Therefore, our restriction digests of the pHAPE plasmid should produce more of the smaller DNA fragments of any particular size than of the larger fragments, thereby increasing the visibility of the lower molecular weight bands. It is also desirable to have “landmark” bands representing particular known lengths significantly brighter than neighbouring bands to facilitate size identification (Table 1). The copy numbers necessary to achieve the desired band intensities were calculated based on the fragment masses seen in commercial ladders using this formula (assuming an average of 650 Da per base-pair):

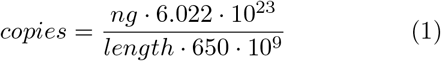

**Table 1:** Fragment lengths from restriction enzyme cleavage of pHAPE. The numbers of fragments of particular lengths produced by cleavages of one plasmid molecule using various restriction enzymes are shown for the HAPE ladder on the left. The ladder produced by only *Bam*HI digestion is on the right. “Proportion” refers to the fraction of the total plasmid mass comprised by fragments of a particular length and can be useful for visual estimation (by comparison) of DNA mass in electrophoresis bands of interest (when the total mass of pHAPE loaded into a gel size marker lane is known). When making such estimations using the HAPE ladder, it is important to consider in calculations the proportion that any restriction digest contributes to the final ladder. Note also that the *Pst*I and *Hind*III digests of pHAPE both produce 1000 bp fragments.

| HAPE |  |  | *Bam*HI |  |  |
| --- | --- | --- | --- | --- | --- |
| Length (bp) | Copies | Proportion | Length (bp) | Copies | Proportion |
| 10000 | 1 | 1.000 | 1178 | 1 | 0.118 |
| 6000 | 1 | 0.600 | 979 | 1 | 0.098 |
| 4000 | 1 | 0.400 | 943 | 1 | 0.094 |
| 3000 | 1 | 0.300 | 800 | 1 | 0.080 |
| 2500 | 1 | 0.250 | 700 | 1 | 0.070 |
| 2000 | 1 | 0.200 | 600 | 1 | 0.060 |
| 1500 | 1 | 0.150 | 500 | 2 | 0.100 |
| 1000 | 1 | 0.100 | 450 | 1 | 0.045 |
| 1000 | 2 | 0.200 | 400 | 1 | 0.040 |
| 950 | 1 | 0.095 | 350 | 1 | 0.035 |
| 800 | 1 | 0.080 | 300 | 2 | 0.060 |
| 700 | 1 | 0.070 | 250 | 1 | 0.025 |
| 600 | 1 | 0.060 | 200 | 5 | 0.100 |
| 500 | 5 | 0.250 | 150 | 2 | 0.030 |
| 400 | 2 | 0.080 | 100 | 2 | 0.020 |
| 300 | 2 | 0.060 | 50 | 5 | 0.025 |
| 200 | 2 | 0.040 |  |  |  |
| 100 | 5 | 0.050 |  |  |  |

In designing the locations of restriction enzyme cleavage sites in pHAPE, fragments of similar sizes were allocated to single enzymes to allow versatility in the combination of the resulting fragments (Figure 1). For example, the *Hind*III digest (which produces the smaller fragments of the HAPE ladder) can produce a ladder in its own right with 100 bp increments up to 1 kb. A final design consideration was to utilise restriction sites for only common, inexpensive restriction enzymes. We chose *Hind*III, *Apa*I, *Pst*I, *Eco*RI, and *Bam*HI. Of course, for single-site cleavage of pHAPE, numerous alternatives to *Apa*I exist in the pBluescript II SK+ derived sequence.

**Figure 1:**
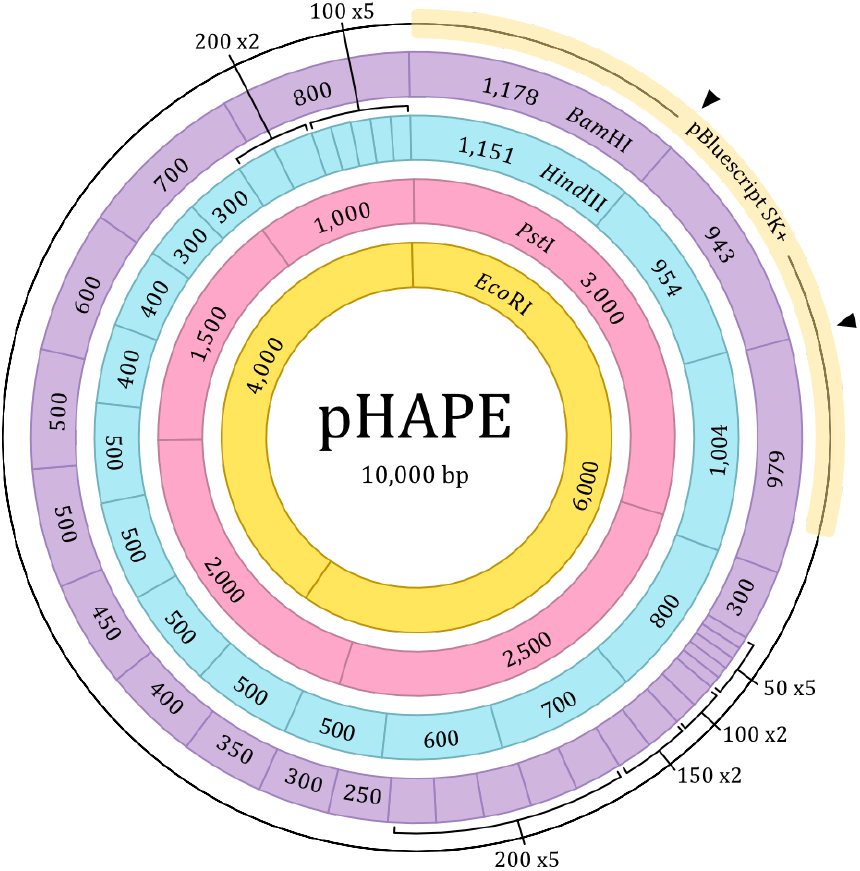
Restriction site map of pHAPE, showing the fragment lengths (in bp) from each restriction enzyme digest. *Eco*RI in yellow, *Pst*I in pink, *Hind*III in blue, and *Bam*HI in purple. The position of the pBluescript II SK+ backbone is annotated, with arrows indicating the positions where *Hind*III and *Bam*HI restriction sites were introduced.

In order to produce all the fragments desired using a minimum number of restriction enzymes, the DNA of the pBluescript II SK+ backbone (2,859 bp) requires modification to include additional *Bam*HI and *Hind*III cleavage sites on either side of the pBluescript II SK+ AmpR gene (Figure 2). To reduce the likelihood of these affecting propagation of the plasmid, we sought to alter as few bases as possible. Ultimately, we devised a scheme where five and four substitutions each are made at two sites flanking AmpR. The modifications are achieved by PCR amplification using mismatched primers (sequences and PCR conditions are provided in the Supplementary Data File) followed by *Dpn*I digestion and Gibson (isothermal) assembly (Gibson, 2009). The introduced *Bam*HI and *Hind*III restriction sites are placed sufficiently close together that single mismatch primers can be used to incorporate the required mutations at each site (Figure 2).

**Figure 2:**
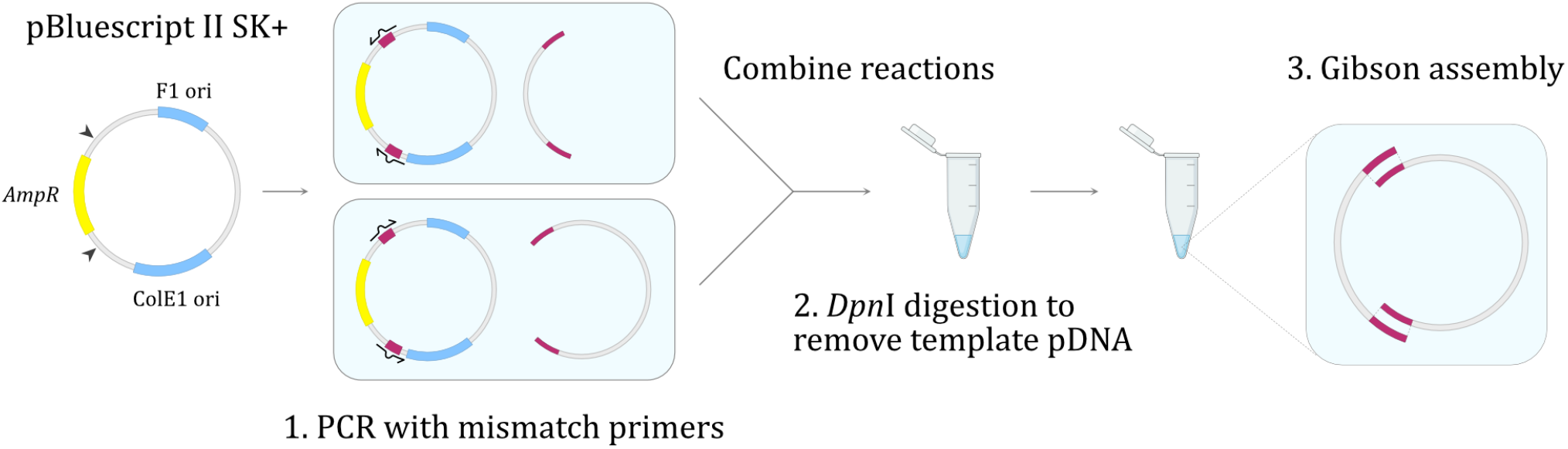
Site-directed mutagenesis of pBluescript II SK+ before insert inclusion to form pHAPE. Mismatch primers were used to introduce two *Bam*HI and *Hind*III sites to the pBluescript II SK+ backbone either side of *AmpR*. Template pDNA was digested with *Dpn*I before Gibson (isothermal) assembly to assemble the modified form of pBluescript II SK+.

### 2.2 Construction of pHAPE

Initially, we attempted to assemble the designed insert sequence by synthesis of three fragments of ~2,500 bp each flanked by unique restriction sites for directional cloning (see Supplementary Data File). To preserve and amplify the synthetic sequences, these were first cloned individually and separately into pBluescript II SK+ to form “Insert Fragment Plasmids”. Despite each insert fragment harbouring restriction sites with unique sticky ends, a four-component simultaneous ligation of the purified fragments with the modified pBluescript II SK+ backbone vector (described in Figure 2) proved unsuccessful. Therefore, we assembled the pHAPE plasmid through a series of sequential ligations and transformations, adding one fragment to the vector at a time, shown in Figure 3.

**Figure 3:**
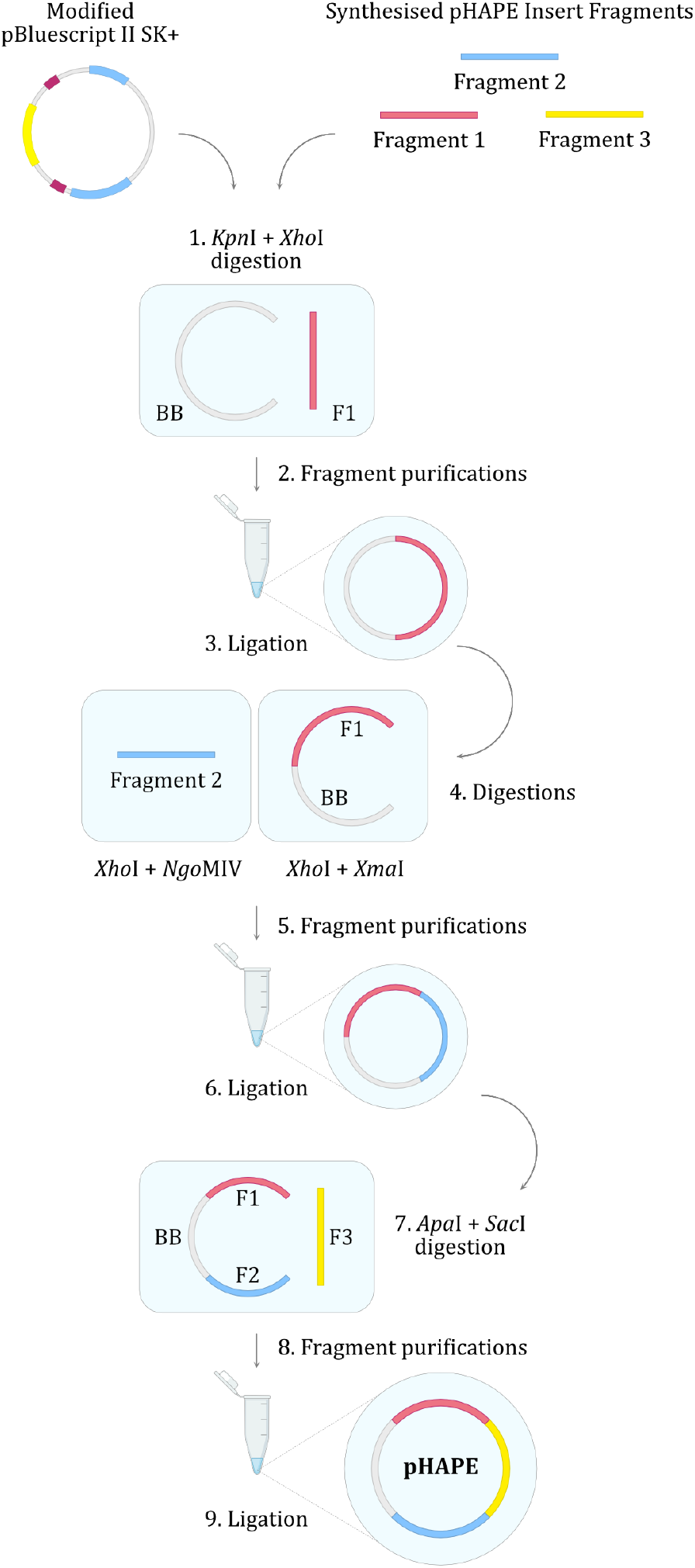
Cloning scheme for assembly of pHAPE. BB - backbone (modified pBluescript II SK+ backbone, see Fig 2), F1 - Insert Fragment 1, F2 - Insert Fragment 2, F3 - Insert Fragment 3.

However, we discovered subsequently that the reason for our lack of success with four-component simultaneous ligation may have been due to plasmid instability caused by one of the fragments (Insert Fragment 1a - see Supplementary Data File). The suspected cause of instability was 13 consecutive *Bam*HI sites intended to produce 25 bp fragments (Lovett et al., 1994; Oliveira et al., 2008). Removal of these sites would not significantly affect the design of the ladder (as the region of DNA allocated to the 25 bp fragments would, instead, form an additional 300 bp fragment). Therefore, a 305 bp section of the sequence was redesigned to abolish those BamHI sites. A new DNA fragment was synthesised and used as a “mega-primer” in site-directed mutagenesis of the Insert Fragment 1a plasmid to form plasmid Insert Fragment 1b (see the Supplementary Data File for the sequence of the mega-primer, the conditions used for mutagenesis, and the intended sequence of Insert Fragment 1b).

When grown side-by-side, colonies harbouring Insert Fragment 1b appeared to grow faster than those with the original sequence (Insert Fragment 1a), and transformation efficiency also appeared to be higher for the Insert Fragment 1b plasmid (data not shown). Note that the *Escherichia coli* strain DH5α was used in all the cloning work above except that strain Stbl3™(Invitrogen, C737303) was used when performing the cloning experiments with Insert Fragment 1a and 1b (due to the relatively closely spaced repetitive restriction sites Insert Fragment 1a contains) as Stbl3™is RecA deficient (*RecA13*). We have found that the assembled pHAPE plasmid can be maintained in *E. coli* DH5α with no apparent instability.

Since it is approximately three times the mass of pBluescript II SK+, the pHAPE plasmid transforms bacteria at a somewhat lower rate than the former plasmid. For example, in one side-by-side comparison, the transformation of chemically competent DH5α cells with pHAPE yielded 5.7e5 cfu/μg of DNA while transformation with pBluescript II SK+ produced 1.434e7 cfu/μg of DNA. Over 100 μg of plasmid DNA can be obtained from a 50 mL culture in Lysogeny Broth (LB) medium containing 100 μg/mL ampicillin. This is sufficient to load >1,000 standard (3 mm) gel lanes with the HAPE ladder and >3,000 lanes with the *Bam*HI ladder.

### 2.3 Ladder assembly

The pHAPE plasmid should be digested separately with *Hin*dIII, *Apa*I, *Pst*I, and *Eco*RI before combination of the digest products to assemble the HAPE ladder with fragments ranging from 10 kb to 100 bp. (The *Apa*I digest can be substituted with any other enzyme that cuts the pHAPE plasmid once.) It should be noted that longer digestion times than typically recommended may be needed, due to the large number of restriction sites present and the supercoiled nature of the purified plasmid (Snounou and Malcolm, 1984). Following digestion, all four enzymes above can be inactivated by heating to 80°C for 20 minutes. Assuming an equal concentration of pHAPE in each digest, these can subsequently be combined in a volume ratio of 20:1:5:2.5 (HindIII, *Apa*I, *Pst*I, *Eco*RI) to produce a visually favourable combination of band intensities. Examples of pHAPE-derived fragment ladders are shown in Figures 4 and 5. Alternatively, the fragment proportions provided in Table 1 can be used to tailor the ladder to the specific needs of a laboratory. To the resulting combination of digests, 1 volume of 6x gel loading dye and 4 volumes of tris-EDTA buffer (10 mM tris, 1 mM EDTA, TE) pH 8 can be added before use without any purification steps. However, the inclusion of a purification step followed by resuspension of the ladder in TE buffer will likely improve its storage longevity and give less distortion of electrophoresis bands due to dissolved salts. Another advisable practise to reduce distortion of DNA fragment “bands” during electrophoresis is to limit the mass of DNA loaded into a gel electrophoresis well to a maximum of 15 ng of DNA per 1 mm^2^ of that surface of the well into which the DNA migrates (~150 ng for 3×3 mm wells).

**Figure 4:**
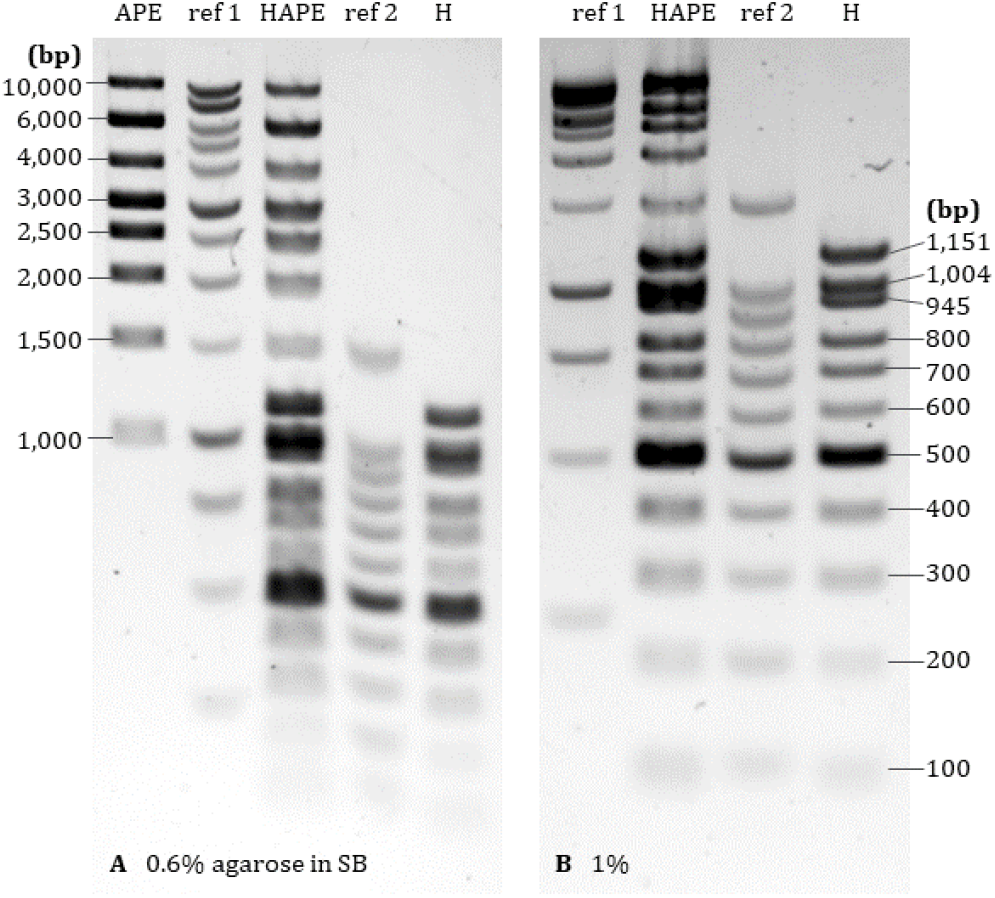
Electrophoresis of pHAPE-derived DNA ladders through 0.6% and 1% agarose gels in SB. Both gels contain 0.5 μg/mL ethidium bromide and were run at 90V for 1 hour. Letters indicate the restriction enzyme digests of pHAPE used to make each ladder: H - *Hind*III, A - *Apa*I, P - *Pst*I, E - *Eco*RI. Reference ladders are the Promega 1 kb (ref 1) and 100 bp (ref 2) ladders.

**Figure 5:**
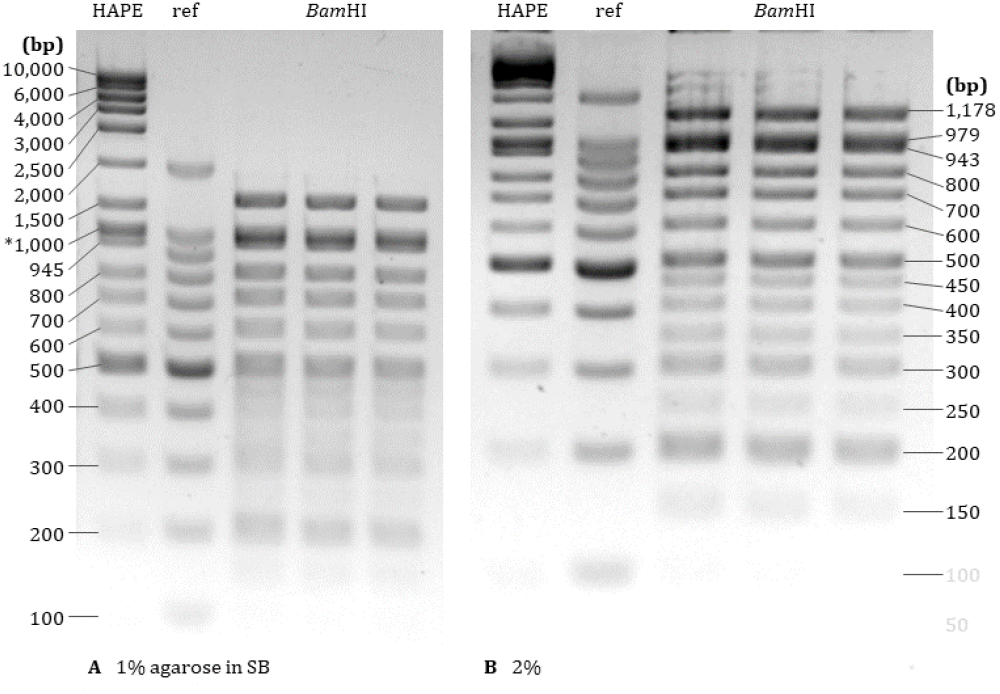
Electrophoresis of the HAPE and *Bam*HI DNA ladders through 1% and 2% agarose gels in SB. Both gels contain 0.5 μg/mL ethidium bromide and were run at 90V for 1 hour. The Promega 100 bp ladder was used as a reference.

### 2.4 Gel Electrophoresis

The HAPE ladder was compared to commercial reference ladders on 0.6-2% agarose gels in sodium borate (SB) buffer in Figures 4 and 5. Fragments of the HAPE ladder were successfully resolved on a 1% gel after 1 hour at 120V.

The pHAPE ”*Bam*HI” ladder (with fragments ranging from 50 bp to 1 kb) is produced with a single *Bam*HI digest. As *Bam*HI cannot easily be inactivated by heating, protein removal (eg. by phenol-chloroform extraction, precipitation using ethanol, and then redissolution in TE buffer) following digestion is recommended. We have not determined whether commercial DNA cleanup kits effectively recover fragments of all lengths from the digests. However, if intending long-term storage of pHAPE digests, we recommend purification and/or frozen storage as small aliquots until needed.

A protocol for the generation of the HAPE ladder from pHAPE can be found in the Supplementary Data File. Alternatively, the fragment proportions provided in Table 1 can be used to tailor ladders to the specific needs of a laboratory.

In summary, the pHAPE plasmid can be generated by mutagenesis of the commonly used pBluescript II SK+ phagemid followed by insertion of 7141 bp (comprised of three smaller fragments). It can be used for generation of a variety of DNA size marker ladders via simple restriction enzyme digestions.

## 3 Materials and Methods

### pHAPE Construction

Protocols for molecular cloning were adapted from (Sambrook and Russell, 2001). Restriction endonucleases, T4 DNA ligase, Gibson assembly enzyme master mix, and all reaction buffers were from New England Biolabs Inc. (Ipswich, Massachusetts, USA). Custom insert fragments for pHAPE and the 305 bp primer used to modify insert fragment 1a were synthesised as IDT gBlocks™ (Integrated DNA Technologies, Inc., Coralville, Iowa, USA). *E. coli* and plasmid DNA were electroporated in 1 mm cuvettes at 1.8 kV using the MicroPulser electroporator (Bio-Rad Laboratories, Inc., Hercules, California, USA). This was performed at room temperature to enhance transformation efficiency (Tu et al., 2016).

### Gel Electrophoresis

Gels were composed of agarose in 1x sodium borate (SB) buffer (36.4 mM boric acid, 10 mM NaOH, pH 8) with 0.5 μg/mL of ethidium bromide. Electrophoresis was performed in 1x SB at 90-120 V for 1 hour. All gels were 6 x 10 cm (width x length) and 4 mm thick, with 3 x 1 x 3 mm (width x length x depth) wells.

## 4 Availability

All sequences required to construct pHAPE can be found in the Supplementary Data File.

We request that any researchers using the pHAPE design acknowledge the work of the authors of this paper by citation.

## 5 Acknowledgements

The authors wish to thank Jordan Pederick for advice on molecular cloning and Gibson assembly, and Dr James Botten for guidance regarding bacterial culture. AA would like to thank Christopher Fusco for helpful discussion and initial drafting of this manuscript. This work was supported by funds from the School of Biological Sciences of the University of Adelaide.

