## Supplementary Data File for "pHAPE: a plasmid for production of DNA size marker ladders for gel electrophoresis"

**pHAPE sequence (10,000 bp)**

Uppercase bases are from the pBluescript II SK+ backbone, lowercase bases are the designed insert. The modified bases in pBluescript have been highlighted blue, and the *AmpR* gene is highlighted green. Insert fragment 2 is highlighted yellow, and is flanked by fragments 1 and 3. Restriction sites used for cloning are underlined: (in order) *Kpn*I, *Xho*I, *Apa*I, *Sac*I.

```
5’-CAGCTTTTGTTCCCTTTAGTGAGGGTTAATTGCGCGCTTGGCGTAATCATGGTCATAGCTGTTTCCTGTGTGAAATTGTT
ATCCGCTCACAATTCCACACAACATACGAGCCGGAAGCATAAAGTGTAAAGCCTGGGGTGCCTAATGAGTGAGCTAACTCACA
TTAATTGCGTTGCGCTCACTGCCCGCTTTCCAGTCGGGAAACCTGTCGTGCCAGCTGCATTAATGAATCGGCCAACGCGCGGG
GAGAGGCGGTTTGCGTATTGGGCGCTCTTCCGCTTCCTCGCTCACTGACTCGCTGCGCTCGGTCGTTCGGCTGCGGCGAGCGG
TATCAGCTCACTCAAAGGCGGTAATACGGTTATCCACAGAATCAGGGGATAACGCAGGAAAGAACATGTGAGCAAAAGGCCAG
CAAAAGGCCAGGAACCGTAAAAAGGCCGCGTTGCTGGCGTTTTTCCATAGGCTCCGCCCCCCTGACGAGCATCACAAAAATCG
ACGCTCAAGTCAGAGGTGGCGAAACCCGACAGGACTATAAAGATACCAGGCGTTTCCCCCTGGAAGCTCCCTCGTGCGCTCTC
CTGTTCCGACCCTGCCGCTTACCGGATACCTGTCCGCCTTTCTCCCTTCGGGAAGCGTGGCGCTTTCTCATAGCTCACGCTGT
AGGTATCTCAGTTCGGTGTAGGTCGTTCGCTCCAAGCTGGGCTGTGTGCACGAACCCCCCGTTCAGCCCGACCGCTGCGCCTT
ATCCGGTAACTATCGTCTTGAGTCCAACCCGGTAAGACACGACTTATCGCCACTGGCAGCAGCCACTGGTAACAGGATTAGCA
GAGCGAGGTATGTAGGCGGTGCTACAGAGTTCTTGAAGTGGTGGCCTAACTACGGCTACACTAGAAGGACAGTATTTGGTATC
TGCGCTCTGCTGAAGCCAGTTACCTTCGGAAAAAGAGTTGGTAGCTCTTGATCCGGCAAACAAACCACCGCTGGTAGCGGTGG
TTTTTTTGTTTGCAAGCAGCAGATTACGCGCAGAAAAAAAGGATCTCAAGAAGATCCTTTGATCTTTTCTACGGGGTCTGACG
CTCAGTGGAACGAAAACTCACGTTAAGGGATTTTGGTCATGAGAGGATCCAAAAGCTTCTTCACCTAGATCCTTTTAAATTAA
AAATGAAGTTTTAAATCAATCTAAAGTATATATGAGTAAACTTGGTCTGACAGTTACCAATGCTTAATCAGTGAGGCACCTAT
CTCAGCGATCTGTCTATTTCGTTCATCCATAGTTGCCTGACTCCCCGTCGTGTAGATAACTACGATACGGGAGGGCTTACCAT
CTGGCCCCAGTGCTGCAATGATACCGCGAGACCCACGCTCACCGGCTCCAGATTTATCAGCAATAAACCAGCCAGCCGGAAGG
GCCGAGCGCAGAAGTGGTCCTGCAACTTTATCCGCCTCCATCCAGTCTATTAATTGTTGCCGGGAAGCTAGAGTAAGTAGTTC
GCCAGTTAATAGTTTGCGCAACGTTGTTGCCATTGCTACAGGCATCGTGGTGTCACGCTCGTCGTTTGGTATGGCTTCATTCA
GCTCCGGTTCCCAACGATCAAGGCGAGTTACATGATCCCCCATGTTGTGCAAAAAAGCGGTTAGCTCCTTCGGTCCTCCGATC
GTTGTCAGAAGTAAGTTGGCCGCAGTGTTATCACTCATGGTTATGGCAGCACTGCATAATTCTCTTACTGTCATGCCATCCGT
AAGATGCTTTTCTGTGACTGGTGAGTACTCAACCAAGTCATTCTGAGAATAGTGTATGCGGCGACCGAGTTGCTCTTGCCCGG
CGTCAATACGGGATAATACCGCGCCACATAGCAGAACTTTAAAAGTGCTCATCATTGGAAAACGTTCTTCGGGGCGAAAACTC
TCAAGGATCTTACCGCTGTTGAGATCCAGTTCGATGTAACCCACTCGTGCACCCAACTGATCTTCAGCATCTTTTACTTTCAC
CAGCGTTTCTGGGTGAGCAAAAACAGGAAGGCAAAATGCCGCAAAAAAGGGAATAAGGGCGACACGGAAATGTTGGATCCTCA
TAAGCTTCCTTTTTCAATATTATTGAAGCATTTATCAGGGTTATTGTCTCATGAGCGGATACATATTTGAATGTATTTAGAAA
AATAAACAAATAGGGGTTCCGCGCACATTTCCCCGAAAAGTGCCACCTAAATTGTAAGCGTTAATATTTTGTTAAAATTCGCG
TTAAATTTTTGTTAAATCAGCTCATTTTTTAACCAATAGGCCGAAATCGGCAAAATCCCTTATAAATCAAAAGAATAGACCGA
GATAGGGTTGAGTGTTGTTCCAGTTTGGAACAAGAGTCCACTATTAAAGAACGTGGACTCCAACGTCAAAGGGCGAAAAACCG
TCTATCAGGGCGATGGCCCACTACGTGAACCATCACCCTAATCAAGTTTTTTGGGGTCGAGGTGCCGTAAAGCACTAAATCGG
AACCCTAAAGGGAGCCCCCGATTTAGAGCTTGACGGGGAAAGCCGGCGAACGTGGCGAGAAAGGAAGGGAAGAAAGCGAAAGG
AGCGGGCGCTAGGGCGCTGGCAAGTGTAGCGGTCACGCTGCGCGTAACCACCACACCCGCCGCGCTTAATGCGCCGCTACAGG
GCGCGTCCCATTCGCCATTCAGGCTGCGCAACTGTTGGGAAGGGCGATCGGTGCGGGCCTCTTCGCTATTACGCCAGCTGGCG
AAAGGGGGATGTGCTGCAAGGCGATTAAGTTGGGTAACGCCAGGGTTTTCCCAGTCACGACGTTGTAAAACGACGGCCAGTGA
GCGCGCGTAATACGACTCACTATAGGGCGAATTGGGTACCctactttctgatggagggtcatcggttcagtgactcgctcgcc
tcagagaactacgcgttgccatgtaatacgctgccagtggtcgcgtgtaggacagggtatatccagttatccccagccagcct
gcaggactatatccaccccgtttcagtcttagtggacacattgcagtctgacccgacggatccgcacctggggtgccttgtag
catgcgccaaagcttccgccaaaatgatccggctggagctagtccgcatggatacatggagatgccagtagcaaggattccac
aactggtggctccgatagatccgacgcggaacgtgggaggagtctccaatgggatgtgcccgtgatgcatacaaagaaaggtc
catcggcctgaccctcctttaattccataacatgatacaaagctcggacttcttaagcgaaccactttccattggtataaat
cgattcccgcgctgctggctaactcggatccatccgtagtgactattgccaattaagttattgcaagtgggagtgggatccaa
cagtgagtgttgcctctggcgtacaccctcctaaaagttaggggatcccatcatactaacactattaaggtttgattattctg
tgcggctaaggatccacatggtccccatgacacgagaccgggaacagaacggggcgtacggatccgaaatagggacgccctct
cccaatgaaaaccttcgcggacagaaggatccaaacacctctcaaatggtttactgtgtaggatgtgtgatgatcaacgcgat
cgttattgtgattcctattgttcctatttcttcagctatatatggatcccaaggattggctgttaattgctgtcttccacttg
gaaccgtccatatacctgagtttcacgcaagcctgatatactcaccttttaaccatccgcggatcctcagctcggattccgtg
caatgtagtagtgggctgtaggacacgatagtcaattggagtttaatgatttatcgaagatgaagcttgtgggtatcgcgttt
gccctacacacgaaattgtttctaggctggtatgcgcatgtagcggatccgccgccctcggttgtggaagaactatagattcc
actgctacttggagcatcacgtatcgtaactagaatcgggtcccgtgggtggccgggtaagcatgagagtaaaacgagtttcc
ctcgcgactgtgacaatgtcgcgcgattggatcctgtgccgtcattcggtggagaagtgaccgcagttccttatcccccccag
ggtatgtcccaaccatgttgtctgggccaagtggctgagtttacgcactggagtggatgcgggcacaaaagtaagcaatacag
gtcctaatatgtggtaaactgcatacagggcctactatattaaagtgggatgtacaaagcgtggatccctttgatagtatgaa
tacagagaaaaatttccgctctgcggtggcatcgctattacctaagtgagtgttcataccgccatggtgtacgttaaagctaa
ctcgactgttgcgcggagaccccttcatttgttagttccaacttgggtccttcacttcttgctatttccagaagtgcctgccc
ataatatactagaggatcctacccattctagcagtgttaccacacccaatacggccagcgggatactgtcattgcacggttct
```

ctgcaatattcttcaaagcttacacattatcagggtactttttgacccccgcgacgtcgattggacaaccggccgcagtcgtc
ttagtactccctcctgttcgtcctcgggcggaacaattcacaagtagggatccccaaaggaagcctcctaaacatttggtcca
ttgtcataccacatttcctagcggaacgtaataaagaagcgtaccacgggggcaagcctcgcaatgtctaatctgtgggatga
gctttaaaggtacgacgtaggagcaagaaacgactagtaggataacgccgtctttgccagcatctccacgttgtcgaatttgg
atccaccataattagttgcgggaacctcgcgcagtcattctgcgaacgcccatgcctaagggatattcccagagtgagcctta
ggtggagtcggcgctaggatgaaaagaatgcattggctgttccagggtgctttgggtgccgccgtggtctgtatttaacgttg
atttggttcgttgaacggggcgcgggcgatgggatccacacgggtgcctgctcaacgctggctgtattctatagacagttca
gaaaatgattagcggtgccttgggggaaagaggaaagcttactgctttgcctgagtactaattcaacctcctatccgatcatg
tcagcggagggcgccacgactgtgttatcgcgctttaagtggtacgaccgcatggttgaacagtaaactgtatgtgttactga
tgagatcgggtgacagattcgcacaatatctgcggatccctctggctactcgagccaacgccgtgttttgctgggagcgggct
tcctcctttagggcgccaaaagcaatgctcggagtggaggcccaacggagcaacaactttcattctccgaggtatggcctcag
tgcgtcatctgcaggtgaattactcgcctatgacgatggcggagtaactgaattatagtatgcggccgccaacggctgctccg
ccaaacagctgctgcgttttcactaatgaaccgaaaagaggccgagagtacgcaaacagccgcgtgagctgcgacttgctgac
tggatccgattattccgggcttaccacgacattaccaagctttagaacgtcatgcatggtcgtcagtagtcaccgaggtaatt
ctaacgcgatccccatgcagcatacaaggtacagggaagccgtcgggccaccgtatggcccgctgctaaagccgttctctgga
tttttctgaaaacataaggctatggagaattgcaaccgccagcgaccaatcagcgcatttaaaatcaggcctaaatcgagtat
attatcgcggtgtttctttggccgcagcagacgtgtcggtcaaattccctcggatcttgagtccgaatcaagacgtcgcgaat
tcccccctaggcgggctctcggatcccatacctcaggagtgtgtccgcccacttgctgttcgcttttaactgtcattgcagccc
cgtagtggactggtggtcaagacccaaccccaatgtttcatcagcacaacttcgatcgacaggtaagcgccaacgtttgcact
gtatactaacgagcactgcgcgggacgaaacgtaggtgaagcttgtttatattagcagaccacccgttgagcccacccttcag
gctgcgaagagtcacggtgtgtaagagaatctgtaaaataatacaggcagagacgctcatgcgtgctggtgcgtggagtaggg
tgagctgcgtttataataatatgttccacctatattgttacttaggcccccaaaactttcgagactccgatatgctttataca
aatgggatccatatctatatgcacctgctacgacccgttttaggagttgagaagtatattccgtagcagcagctctagcttct
acctccaatttataaccctgcggttccggtcatcgcatgggggtcaacggagggccacagtcaacagtgaagtagcgccttga
cgcccttatacagtttgcgagttaggtgaggatagtcttcccatatgggtgttgatcaggtagacaaagttgtaggcaggcag
cagggcaccgaactttccgactctgcgagcacatataagcaagcttactgctgacctgtctggcataggaaggaaactgctat
aattctgagggcggcccctatatcgtggtaagttcggggatagcactgcgaggaaattcgttcaagtccgtcaagattgattg
tgctaaaaggattcatatcactggttgatacctcatcctggatccacccgggtccaagggtacgccagcccatttaatgagcc
gaatactttcacggctcaaatcgggatgcctggccccgactatgtgaaataaacacactatccgtatggtctacggggcagct
ggtaaccaacgtgtgatacactgtacgggtgtccgccgtgatcacaacttgcccgtcatgtgaggctacgaataatagctccg
gtactacaaccactgtagtcgtgagagcctgggccgtggcacgtaccacggaagattaaatgtttcctcacgcgtcagaagtt
cactgaggtaacgagcattgactatgaagtagttaggtcggtaagcttcattacccagcttcgaccactttgcactacaccct
agatgaagcactctttcgtacgtcgtaagttcctcttgaagtccgagtggcaccctgccaaaacgtccgacggggttcctgca
tgttatgttcctaatgaaggtacggccaagctcgcacgcgaggatccctcgtagaggctaccggacacgtatctgcgtattag
ggatcgggtcacggagccgcgtcaccacattggagtgaatacgcacaataaacgccgggtgcagaatccttctatgcatcgtg
tattgagtaaatgcttctgcagaccccaacgcttcactccggatatggcagtaagataataggtatccgctgctaaaaggcgt
cgagatcagccagcacactataatgcataacggccttaaacctctcgggactcatagagattgcttcagcaaactgtcaagtc
cgtccacgaatgcgcaattgtgtaacattcgcccggccatccagaagcttgaaggattagccctctgaacccagctgttcaga
ggccgtaatagtctgcggtctctggcgtggtatatttatatgatctctagctacgtttgtcttctggtcaacaatacgagaca
gttggagcgggggctcgcaatgggaaaaggacatcggcctaatggatcctaaccgggccccgagtccaacagaacatgccgag
cgggggcaggtagcgttcaattcacatgtgagcacactgttggcgattaccctggatttcatgcatctttggagcggaccgg
agcattggttctctaaaccgagcgaggcggtcggaggtcaataaaggtgcgggtcacgcaaagaatagatccgtgtggtcgat
caagcgtgacatactgcgcacgccggtcgaagcttgcggccgtctacctttcgcgtcattagttattacctcacccacccagt
cccatgttgagttttatgatgaccgagcccccgttccagcccggccaaaatcattatcagtctgactaattgcggcaactttc
aatatccccgatcagctagttccaaatcagggactcacacgggtatcattgttgcctggatcttgtggacggtcaacacctaa
tggtaagaaaggtgactcaggtcttaaccggtatctgagcaggtttctaaggatgaacgaatttgctgatcccattgttcgcg
tgacagcccgcacacgtatgtgaaaccacgttgcggagcatggtgagattaacagaacgggggatccggtatactcatgcga
gcaccaaactagcaaagcttagaacgattttgttattaagcagtgacttacgtatagcttccgcccagtccactgtttgtgat
tcatcagcagatgggactaggtaatggcactgatcgtccgggtgaggtatataggtaacaggggtattcgcttcacaaaataa
ctgcgggtatccttcgtgctcatactcgatgcacctgccccacattggattacgagatccgttccacatgtaccgtcatacga
gctgacccagacatcagcaccgtaatcctgatttggaacgtaaagggttatggcggcggacaaaaaagcttagttacgctgta
tcgctttgtccgcataattcatgcctcctatgatgagcgcgccagtatattacgtgttagccgaactccatttgtcaccagta
attcgcaagcacagtgggaatcccgaggaatatatgtcttccattgaacgacccgttgctagacgggggtcacccagtatgac
agcttgagagaagagacacgcactgcagttagtacggtaggagtaaacagtctatgcgcttgaccctgaattaggctgtcgtg
gttaacagtctccgtggttggaagtttacagccaagcttacatgacataagacgagtgctaaggctctttttttaaccaggttc
ccttaatgactacgcggatccccagtgctaaaaactcgtgtgccgtcgagggcaacgtgtgcccgtcatgggcgcaccacgct
ccaagtacgtattaaactgtgactcaggatggagagacctgcccctagtctaacgaaaaacagagtgaagcttgatctcattt
gacgccccgaagtgagatgattccttgctacccagaagtttgtggcccatgaaagtccttctggcctaccttcgacaagatcg
tccagtaaaaattgtgtcatatggacgataaatcggctttaacaaagatccacacacgtctatcggactcctaacgttgaagg
actcttgcgcacgagccgaagcttctcacccacctgagccctacctaagctgagctatacgttcacttcctgcggacatgacg
aggaccagttcaacggggacggtataatagccatgaagcttaccaccagtctggagccgctctgatggctcgaccaaccttta
tcgtcaccactttaaaccaacagatcaattggtctttcccaagcgagcacgaaagcttgctactaccccactggtggttcggg
tcgttgaaatacgccttcttccttatttgctgtacgcttgaaacgagtgatggtagcctgagtcgggggaagcttatcgaagg

cataggcgaggcgtaatacatgcgatcctcgcagcattgcagcggagccagactctattccgaacttaaaccactcataaata
agcaagctttgaactaatccctcgaaatgtgtgtgttcattaatgaaacccatctcagagaaactcgcggatccataaggcgc
gataatatgcttgaattctcaagcttctgcagctaaggagctC-3’

#### Insert Fragment 1(b) sequence

This sequence contains the flanking restriction sites and overhanging bases of the original Insert Fragment 1a sequence, but with the internal sequence of Insert Fragment 1b (*Bam*HI sites abolished).

aaggatcattggtaccctactttctgatggagggtcatcggttcagtgactcgctcgcctcagagaactacgcgttgccatgt
aatacgctgccagtggtcgcgtgtaggacagggtatatccagttatccccagccagcctgcaggactatatccaccccgtttc
agtcttagtggacacattgcagtctgacccgacggatccgcacctggggtgccttgtagcatgcgccaaagcttccgccaaaa
tgatccggctggagctagtccgcatggatacatggagatgccagtagcaaggattccacaactggtggctccgatagatccga
cgcggaacgtgggaggagtctccaatgggatgtgcccgtgatgcatacaaagaaaggtccatcggcctgaccctccttttaat
tccataacatgatacaaagctcggacttcttaagcgaaccactttccattggtataaatcgattcccgcgctgctggctaact
cggatccatccgtagtgactattgccaattaagttattgcaagtgggagtgggatccaacagtgagtgttgcctctggcgtac
accctcctaaaagttaggggatcccatcatactaacactattaaggtttgattattctgtgcggctaaggatccacatggtcc
ccatgacacgagaccgggaacagaacggggcgtacggatccgaaatagggacgccctctcccaatgaaaaccttcgcggacag
aaggatccaaacacctctcaaatggtttactgtgtaggatgtgtgatgatcaacgcgatcgttattgtgattcctattgttcc
tatttcttcagctatatatggatcccaaggattggctgttaattgctgtcttccacttggaaccgtccatatacctgagtttc
acgcaagcctgatatactcaccttttaaccatccgcggatcctcagctcggattccgtgcaatgtagtagtgggctgtaggac
acgatagtcaattggagtttaatgatttatcgaagatgaagcttgtgggtatcgcgtttgccctacacacgaaattgtttcta
ggctggtatgcgcatgtagcggatccgccgccctcggttgtggaagaactatagattccactgctacttggagcatcacgtat
cgtaactagaatcgggtcccgtgggtggccgggtaagcatgagagtaaaacgagtttccctcgcgactgtgacaatgtcgcgc
gattggatcctgtgccgtcattcggtggagaagtgaccgcagttccttatccccccagggtatgtcccaaccatgttgtctg
ggccaagtggctgagtttacgcactggagtggatgcgggcacaaaagtaagcaatacaggtcctaatatgtggtaaactgcat
acagggcctactatattaaagtgggatgtacaaagcgtggatccctttgatagtatgaatacagagaaaaatttccgctctgc
ggtggcatcgctattacctaagtgagtgttcataccgccatggtgtacgttaaagctaactcgactgttgcgcggagacccct
tcatttgttagttccaacttgggtccttcacttcttgctatttccagaagtgcctgcccataatatactagaggatcctaccc
attctagcagtgttaccacacccaatacggccagcgggatactgtcattgcacggttctctgcaatattcttcaaagcttaca
cattatcagggtactttttgacccccgcgacgtcgattggacaaccggccgcagtcgtcttagtactccctcctgttcgtcct
cgggcggaacaattcacaagtagggatccccaaaggaagcctcctaaacatttggtccattgtcataccacatttcctagcgg
aacgtaataaagaagcgtaccacgggggcaagcctcgcaatgtctaatctgtgggatgagctttaaaggtacgacgtaggagc
aagaaacgactagtaggataacgccgtctttgccagcatctccacgttgtcgaatttggatccaccataattagttgcgggaa
cctcgcgcagtcattctgcgaacgcccatgcctaagggatattcccagagtgagccttaggtggagtcggcgctaggatgaaa
agaatgcattggctgttccagggtgctttgggtgccgccgtggtctgtatttaacgttgatttggttcgttgaacgggggcgc
gggcgatgggatccacacgggtgcctgctcaacgctggctgtattctatagacagttcagaaaatgattagcggtgccttggg
ggaaagaggaaagcttactgctttgcctgagtactaattcaacctcctatccgatcatgtcagcggagggcgccacgactgtg
ttatcgcgctttaagtggtacgaccgcatggttgaacagtaaactgtatgtgttactgatgagatcgggtgacagattcgcac
aatatctgcggatccctctggctactcgagcgcaagtggc

#### Insert Fragment 2 sequence

cctggacattctcgagccaacgccgtgttttgctgggagcgggcttcctcctttagggcgccaaaagcaatgctcggagtgga
ggcccaacggagcaacaactttcattctccgaggtatggcctcagtgcgtcatctgcaggtgaattactcgcctatgacgatg
gcggagtaactgaattatagtatgcggccgccaacggctgctccgccaaacagctgctgcgttttcactaatgaaccgaaaag
aggccgagagtacgcaaacagccgcgtgagctgcgacttgctgactggatccgattattccgggcttaccacgacattaccaa
gctttagaacgtcatgcatggtcgtcagtagtcaccgaggtaattctaacgcgatccccatgcagcatacaaggtacagggaa
gccgtcgggccaccgtatggcccgctgctaaagccgttctctggatttttctgaaaacataaggctatggagaattgcaaccg
ccagcgaccaatcagcgcatttaaaatcaggcctaaatcgagtatattatcgcggtgtttctttggccgcagcagacgtgtcg
gtcaaattccctcggatcttgagtccgaatcaagacgtcgcgaattcccccctaggcgggctctcggatcccatacctcaggag
tgtgtccgcccacttgctgttcgcttttaactgtcattgcagccccgtagtggactggtggtcaagacccaaccccaatgttt
catcagcacaacttcgatcgacaggtaagcgccaacgtttgcactgtatactaacgagcactgcgcgggacgaaacgtaggtg
aagcttgtttatattagcagaccacccgttgagcccacccttcaggctgcgaagagtcacggtgtgtaagagaatctgtaaaa
taatacaggcagagacgctcatgcgtgctggtgcgtggagtagggtgagctgcgtttataataatatgttccacctatattgt
tacttaggccccccaaaactttcgagactccgatatgctttatacaaatgggatccatatctatatgcacctgctacgacccgt
tttaggagttgagaagtatattccgtagcagcagctctagcttctacctccaatttataaccctgcggttccggtcatcgcat
gggggtcaacggagggccacagtcaacagtgaagtagcgccttgacgcccttatacagtttgcgagttaggtgaggatagtct
tcccatatgggtgttgatcaggtagacaaagttgtaggcaggcagcagggcaccgaactttccgactctgcgagcacatataa
gcaagcttactgctgacctgtctggcataggaaggaaactgctataattctgagggcggcccctatatcgtggtaagttcggg
gatagcactgcgaggaaattcgttcaagtccgtcaagattgattgtgctaaaaggattcatatcactggttgatacctcatcc
tggatccacccgggtccaagggtacgccagcccatttaatgagccgaatactttcacggctcaaatcgggatgcctggccccg

actatgtgaaataaacacactatccgtatggtctacggggcagctggtaaccaacgtgtgatacactgtacgggtgtccgccg
tgatcacaacttgcccgtcatgtgaggctacgaataatagctccggtactacaaccactgtagtcgtgagagcctgggccgtg
gcacgtaccacggaagattaaatgtttcctcacgcgtcagaagttcactgaggtaacgagcattgactatgaagtagttaggt
cggtaagcttcattacccagcttcgaccactttgcactacaccctagatgaagcactctttcgtacgtcgtaagttcctcttg
aagtccgagtggcaccctgccaaaacgtccgacggggttcctgcatgttatgttcctaatgaaggtacggccaagctcgcacg
cgaggatccctcgtagaggctaccggacacgtatctgcgtattagggatcgggtcacggagccgcgtcaccacattggagtga
atacgcacaataaacgccgggtgcagaatccttctatgcatcgtgtattgagtaaatgcttctgcagaccccaacgcttcact
ccggatatggcagtaagataataggtatccgctgctaaaaggcgtcgagatcagccagcacactataatgcataacggcctta
aacctctcgggactcatagagattgcttcagcaaactgtcaagtccgtccacgaatgcgcaattgtgtaacattcgcccggcc
atccagaagcttgaaggattagccctctgaacccagctgttcagaggccgtaatagtctgcggtctctggcgtggtatattta
tatgatctctagctacgtttgtcttctggtcaacaatacgagacagttggagcgggggctcgcaatgggaaaggacatcggc
ctaatggatcctaaccgggccctgagtccgtt

### Insert Fragment 3 sequence

actcagatgtgggccccgagtccaacagaacatgccgagcgggggcaggtagcgttcaattcacatgtgagcacactgttgg
cgattaccctggatttcatgcatctttggagcggaccggagcattggttctctaaaccgagcgaggcggtcggaggtcaataa
aggtgcgggtcacgcaaagaatagatccgtgtggtcgatcaagcgtgacatactgcgcacgccggtcgaagcttgcggccgtc
tacctttcgcgtcattagttattacctcacccacccagtcccatgttgagttttatgatgaccgagccccccgttccagcccgg
ccaaaatcattatcagtctgactaattgcggcaactttcaatatccccgatcagctagttccaaatcagggactcacacgggt
atcattgttgcctggatcttgtggacggtcaacacctaatggtaagaaaggtgactcaggtcttaaccggtatctgagcaggt
ttctaaggatgaacgaatttgctgatcccattgttcgcgtgacagcccgcacacgtatgtgaaaccacgttgcggagcatggt
gagattaacagaacgggggggatccggtatactcatgcgagcaccaaactagcaaagcttagaacgatttgttattaagcagt
gacttacgtatagcttccgcccagtccactgtttgtgattcatcagcagatgggactaggtaatggcactgatcgtccgggtg
aggtatataggtaacaggggtattcgcttcacaaaataactgcgggtatccttcgtgctcatactcgatgcacctgccccaca
ttggattacgagatccgttccacatgtaccgtcatacgagctgacccagacatcagcaccgtaatcctgatttggaacgtaaa
gggttatggcggcggacaaaaaagcttagttacgctgtatcgctttgtccgcataattcatgcctcctatgatgagcgcgcca
gtatattacgtgttagccgaactccatttgtcaccagtaattcgcaagcacagtgggaatcccgaggaatatatgtcttccat
tgaacgacccgttgctagacgggggtcacccagtatgacagcttgagagaagagacacgcactgcagttagtacggtaggagt
aaacagtctatgcgcttgaccctgaattaggctgtcgtggttaacagtctccgtggttggaagtttacagccaagcttacatg
acataagacgagtgctaaggctctttttttaaccaggttcccttaatgactacgcggatccccagtgctaaaaactcgtgtgcc
gtcgagggcaacgtgtgcccgtcatgggcgcaccacgctccaagtacgtattaaactgtgactcaggatggagagacctgccc
ctagtctaacgaaaaacagagtgaagcttgatctcatttgacgccccgaagtgagatgattccttgctacccagaagtttgtg
gcccatgaaagtccttctggcctaccttcgacaagatcgtccagtaaaaattgtgtcatatggacgataaatcggctttaaca
aagatccacacacgtctatcggactcctaacgttgaaggactcttgcgcacgagccgaagcttctcacccacctgagccctac
ctaagctgagctatacgttcacttcctgcggacatgacgaggaccagttcaacggggacggtataatagccatgaagcttacc
accagtctggagccgctctgatggctcgaccaacctttatcgtcaccactttaaaccaacagatcaattggtctttcccaagc
gagcacgaaagcttgctactaccccactggtggttcgggtcgttgaaatacgccttcttccttatttgctgtacgcttgaaac
gagtgatggtagcctgagtcgggggaagcttatcgaaggcataggcgaggcgtaatacatgcgatcctcgcagcattgcagcg
gagccagactctattccgaacttaaaccactcataaataagcaagctttgaactaatccctcgaaatgtgtgtgttcattaat
gaaacccatctcagagaaactcgcggatccataaggcgcgataatatgcttgaattctcaagcttctgcagctaaggagctcc
gccaaagct

### Insert Fragment 1a (intended sequence)

The region modified in insert fragment 1b is highlighted yellow.

5’-cctactttctgatggagggtcatcggttcagtgactcgctcgcctcagagaactacgcgttgccatgtaatacgctgcca
gtggtcgcgtgtaggacagggtatatccagttatccccagccagcctgcaggactatatccaccccgtttcagtcttagtgga
cacattgcagtctgacccgacggatccgcacctggggtgccttgtaggatccgccaaagcttccgccaaaggatccggctgg
agctagtccgcatggatccatggagatgccagtagcaaggatcccacaactggtggctccgatggatccgacgcggaacgtgg
gaggaggatccaatgggatgtgcccgtgatggatccaaagaaaggtccatcggccggatcctcctttaattccataacagga
tccaaagctcggacttcttaagggatccactttccattggtataaatggatccccgcgctgctggctaactcggatccatccg
tagtgactattgccaattaagttattgcaagtgggagtgggatccaacagtgagtgttgcctctggcgtacaccctcctaaaa
gttaggggatcccatcatactaacactattaaggtttgattattctgtgcggctaaggatccacatggtccccatgacacgag
accgggaacagaacggggcgtacggatccgaaatagggacgccctctcccaatgaaaaccttcgcggacagaaggatccaaac
acctctcaaatggtttactgtgtaggatgtgtgatgatcaacgcgatcgttattgtgattcctattgttcctatttcttcagc
tatatatggatcccaaggattggctgttaattgctgtcttccacttggaaccgtccatatacctgagtttcacgcaagcctga
tatactcacctttaaccatccgcggatcctcagctcggattccgtgcaatgtagtagtgggctgtaggacacgatagtcaat
tggagtttaatgatttatcgaagatgaagcttgtgggtatcgcgtttgccctacacacgaaattgtttctaggctggtatgcg
catgtagcggatccgccgccctcggttgtggaagaactatagattccactgctacttggagcatcacgtatcgtaactagaat
cgggtcccgtgggtggccgggtaagcatgagagtaaaacgagtttccctcgcgactgtgacaatgtcgcgcgattggatcctg
tgccgtcattcggtggagaagtgaccgcagttccttatccccccagggtatgtcccaaccatgttgtctgggccaagtggct

gagtttacgcactggagtggatgcgggcacaaaagtaagcaatacaggtcctaatatgtggtaaactgcatacagggcctact
atattaaagtgggatgtacaaagcgtggatccctttgatagtatgaatacagagaaaaatttccgctctgcggtggcatcgct
attacctaagtgagtgttcataccgccatggtgtacgttaaagctaactcgactgttgcgcggagaccccttcatttgttagt
tccaacttgggtccttcacttcttgctatttccagaagtgcctgcccataatatactagaggatcctacccattctagcagtg
ttaccacacccaatacggccagcgggatactgtcattgcacggttctctgcaatattcttcaaagcttacacattatcagggt
actttttgacccccgcgacgtcgattggacaaccggccgcagtcgtcttagtactccctcctgttcgtcctcgggcggaacaa
ttcacaagtagggatccccaaaggaagcctcctaaacatttggtccattgtcataccacatttcctagcggaacgtaataaag
aagcgtaccacgggggcaagcctcgcaatgtctaatctgtgggatgagctttaaaggtacgacgtaggagcaagaaacgacta
gtaggataacgccgtctttgccagcatctccacgttgtcgaatttggatccaccataattagttgcgggaacctcgcgcagtc
attctgcgaacgcccatgcctaagggatattcccagagtgagccttaggtggagtcggcgctaggatgaaaagaatgcattgg
ctgttccagggtgctttgggtgccgccgtggtctgtatttaacgttgatttggttcgttgaacgggggcgcgggcgatgggat
ccacacgggtgcctgctcaacgctggctgtattctatagacagttcagaaaatgattagcggtgccttggggggaaagaggaaa
gcttactgctttgcctgagtactaattcaacctcctatccgatcatgtcagcggagggcgccacgactgtgttatcgcgcttt
aagtggtacgaccgcatggttgaacagtaaactgtatgtgttactgatgagatcgggtgacagattcgcacaatatctgcgga
tccctctggctac-3’

**Site-directed mutagenesis of fragment 1a**

Mega-primer sequence (305 bp):

5’-gcacctggggtgccttgtagcatgcgccaaagcttccgccaaaatgatccggctggagctagtccgcatggatacatgga
gatgccagtagcaaggattccacaactggtggctccgatagatccgacgcggaacgtgggaggagtctccaatgggatgtgcc
cgtgatgcatacaaagaaaggtccatcggcctgaccctccttttaattccataacatgatacaaagctcggacttcttaagcg
aaccactttccattggtataaatcgattcccgcgctgctggctaactcggatccatccg-3’

PCR conditions:

*Reaction contents (final concentrations)*

| | | |
|---|---|---|
| Insert fragment 1a plasmid DNA | 10 fmol | |
| Mega-primer | 100 fmol | |
| dNTPs | 0.2 mM | |
| Phusion™ HF Buffer | (gives 1.5 mM $MgCl_2$) | (Thermo Scientific F530S) |
| Phusion™ HF DNA Polymerase | 0.02 units | (Thermo Scientific F530S) |

*Cycling conditions*

95°C 2 min, then 20 cycles of:
95°C 30 sec,
70°C 1 min,
72°C 4 min,
and then 72°C 10 min

**Site-directed mutagenesis of pBluescript II SK+**

Mismatched bases are bold and underlined.

Primer pair 1:
Site 1 F 5`-GGGATTTTGGTCATGAGA**GG**ATC**C**AAAAG**CT**TCTTCACCTAGATCCTT-3`
Site 2 R 5`-CAATAATATTGAAAAAGGAAG**CT**TATGAG**G**AT**C**CAACATTTCCGTGTCG-3`

Primer pair 2:
Site 2 F – 5`-CGACACGGAAATGTTG**G**AT**C**CTCATA**AG**CTTCCTTTTTCAATATTATTG-3`
Site 1 R – 5`-AAGGATCTAGGTGAAGA**AG**CTTTT**G**GAT**CC**TCTCATGACCAAAATCCC-3`

PCR conditions:

*Reaction contents (final concentrations)*

| | | |
|---|---|---|
| pBluescript II SK+ (with gene insert) | 2.215 fmol | |
| Herculase II reaction buffer | 1x (gives 2 mM $MgCl_2$) | (Agilent 600675) |
| Forward/reverse primers | 0.5 µM ea. | |
| dNTPs | 0.2 mM | |
| Herculase II Fusion DNA Polymerase* | 0.4 µL in 20 µL reaction | (Agilent 600675) |

*(concentration not provided)

*Cycling conditions*

95°C 2 min, then 30 cycles of:
95°C 30 sec,
60°C 30 sec,
72°C 4 min,
and then 72°C 5 min

Ramp rate = 2°C/sec for all PCRs

**HAPE ladder assembly protocol**

**MATERIALS**

pHAPE plasmid DNA
Water (molecular biology grade)
Restriction endonucleases – *Apa*I, *Eco*RI, *Pst*I, *Hin*dIII
Restriction endonuclease buffers
1x tris-EDTA (TE) pH 8.0

**METHOD**

1. Transfer the following mass of DNA to sterile reaction tubes:

| Reaction | pHAPE (ng) | Final mass ratio in HAPE ladder |
|---|---|---|
| 1 (*Apa*I) | 40 | 1 |
| 2 (*Eco*RI) | 100 | 2.5 |
| 3 (*Pst*I) | 200 | 5 |
| 4 (*Hin*dIII) | 800 | 20 |

2. Set up 20 µL reactions containing the appropriate concentrations of restriction endonuclease and reaction buffer.
3. Incubate the reactions at 37°C (or appropriate digestion temperature for the enzyme used) for 30-60 minutes. Inactivate the reactions by heating to 80°C for 20 minutes.
4. Transfer all reactions to a single 1.5 mL capped tube.
5. (Optional) DNA purification
6. Add 20 µL each of 1x TE (pH 8.0) and 6x gel loading dye (0.25% (w/v) bromophenol blue, 0.25% (w/v) xylene cyanol, 30% (w/v) glycerol[1]). Mix well by gently pipetting up and down.

We recommend that 5 µL of HAPE ladder is loaded per well (for wells 3 x 1 x 3 mm (width x length x depth)).

[1] DNA loading buffer (6X). (2007). Cold Spring Harbor Protocols, 2007(6), pdb.rec11045. https://doi.org/10.1101/pdb.rec11045
